# Geometrically-derived action potential markers for model development: a principled approach?

**DOI:** 10.1101/2024.09.06.611658

**Authors:** Michael Clerx, Gary R Mirams

## Abstract

We explore whether action potential biomarkers can be rederived as functions of a linearisation of the action potential.

## 1. Introduction

Action potential (AP) biomarkers such as “AP duration at 90% repolarisation” (APD90) or “minimum diastolic potential” (MDP) are used to quantitatively compare the shapes of APs recorded from excitable cells [1]. They are also used in modelling, for example as outputs in a sensitivity analysis [2], or as part of an error or likelihood function in model calibration [3].

Central to such applications is the idea that biomarkers capture the essential features of an AP. In this study, we sought to formalise this notion by treating biomarkers as a form of *signal decomposition*. Specifically, we constructed linear approximations of AP waveforms, and then defined biomarkers in terms of these approximations.

Secondly, we aimed to make our biomarkers suitable for use as part of an error function for model calibration. Recent work has shown that using complex algorithms inside an error function can lead to inefficient methods and inaccurate results, and so we tried to make our biomarker calculation as simple as possible by basing it on geometrical methods.

In the next section, we provide an outline of our approach, with results for nodal, atrial, and ventricular AP shapes shown in the Results section.

## 2. Methods

Source code for this study is available at https://github.com/CardiacModelling/GeometricBiomarkers

### 2.1. Data generating and pre-processing

Although our method was developed using experimental data, here we demonstrate it on simulated data generated by a nodal [4], atrial [5], and ventricular [6, 7] cell model. These were chosen to represent three archetypal AP shapes: a self-exciting and relatively slowly depolarising AP, a triangular AP, and an AP with a pronounced spike-and-dome morphology. To mimic experimental data, simulations were “sampled” every 0.1ms, Gaussian noise was added with *σ* = 0.5mV. Next, the data was filtered and downsampled so that each 8 points were replaced by their average (essentially an *n* = 3 Haar wavelet transform).

### 2.2. Geometrical methods

We define two “geometrical” methods which we will use to construct a linear approximation of the AP. **peaks**(*t, v, v*_*t*_, *n*): Finds peaks in the signal. Given a time series (*t, v*), this function (1) isolates all segments where *v* is above the threshold *v*_*t*_ for more than *n* consecutive samples; and (2) returns the time-series indices of the maximum value on each segment.

**rpeaks**(*t*_*s*_, *v*_*s*_, *θ*): Given a time series segment (*t*_*s*_, *v*_*s*_), this method rotates the signal (see below) by *θ* degrees, before identifying the maximum value on the rotated signal and returning its time-series index. To define the concept of a “rotation” in a two-dimensional time-voltage space, we need to define a characteristic scale to normalise each dimension dimension by (“how much time equals how much voltage?”). Based on a typical 1Hz AP going from approximately -90mV to 30mV, we chose 1000ms and 120mV as the characteristic scales.

### 2.3. Linearisation and biomarkers

To construct a linear approximation for a particular type of AP, we start by identifying a “prototypical” signal, and manually performing the following steps: (1) We use the **peaks** method with *v*_*t*_ = −20mV and *n* = 20 (25ms) to identify the peaks in our (downsampled) signal. In all biomarker sets, these peaks function as our first points “P”, and are used for an initial segmentation of the signal into individual APs (where each AP starts and ends as a peak — later we can revise this to start and end APs at their upstroke). (2) We then call **peaks** on the inverse voltage signal with *v*_*t*_ = 0mV and *n* = 20 to find the minimum diastolic point “MDP” for each AP. (3) The remaining approximation is created by drawing lines between points, starting with P and MDP, and either halting if the line is a reasonable approximation, or adding more points. (4a) If the line crosses the AP, we use **rpeaks** to rotate the signal so that the line is horizontal and add two new points representing the minimum and maximum of the rotated signal. (4b) If the line does not cross the AP, we again use **rpeaks** to rotate the signal but this time add only a single new point at the maximum/minimum value above/below the line.

This procedure is followed manually for the prototype signal, and is halted when the approximation is deemed good enough by the designer. Calculation on subsequent signals follows the predetermined procedure designed on the prototype.

The final set of biomarkers is calculated as a function of points derived from the linearisation. For example, the cycle length can be determined as the time from P_i_ to P_i+1_, while the APD might be identified as the time between points indicating the start of the upstroke and full repolarisation.

## 3. Results

We first tried using the methods sketched above to define biomarkers for a self-exciting cell, using as a prototype the simulated data from a sino-atrial node (SAN) model [4]. The derivation procedure is shown in Figure 1. In the signal shown, the peak indicates the peak of the plateau (the weaker sodium current in SAN means there is no strong spike) and the MDP corresponds clearly to the end of the AP. This means we can already derive an “AP magnitude” biomarker from the difference in these voltages. The line from P to MDP intersects the AP, leading to the identification of two points, AP1 and AP2, that can be used to derive quantities with a similar function as the common APD_50_ and APD_90_ biomarkers (although numerically closer to an APD_20_ and APD_95_, respectively). The minimum under the line from MDP to subsequent P identifies a point TO which provides both a take-off potential and a clear start to the following AP.

**Figure 1.**
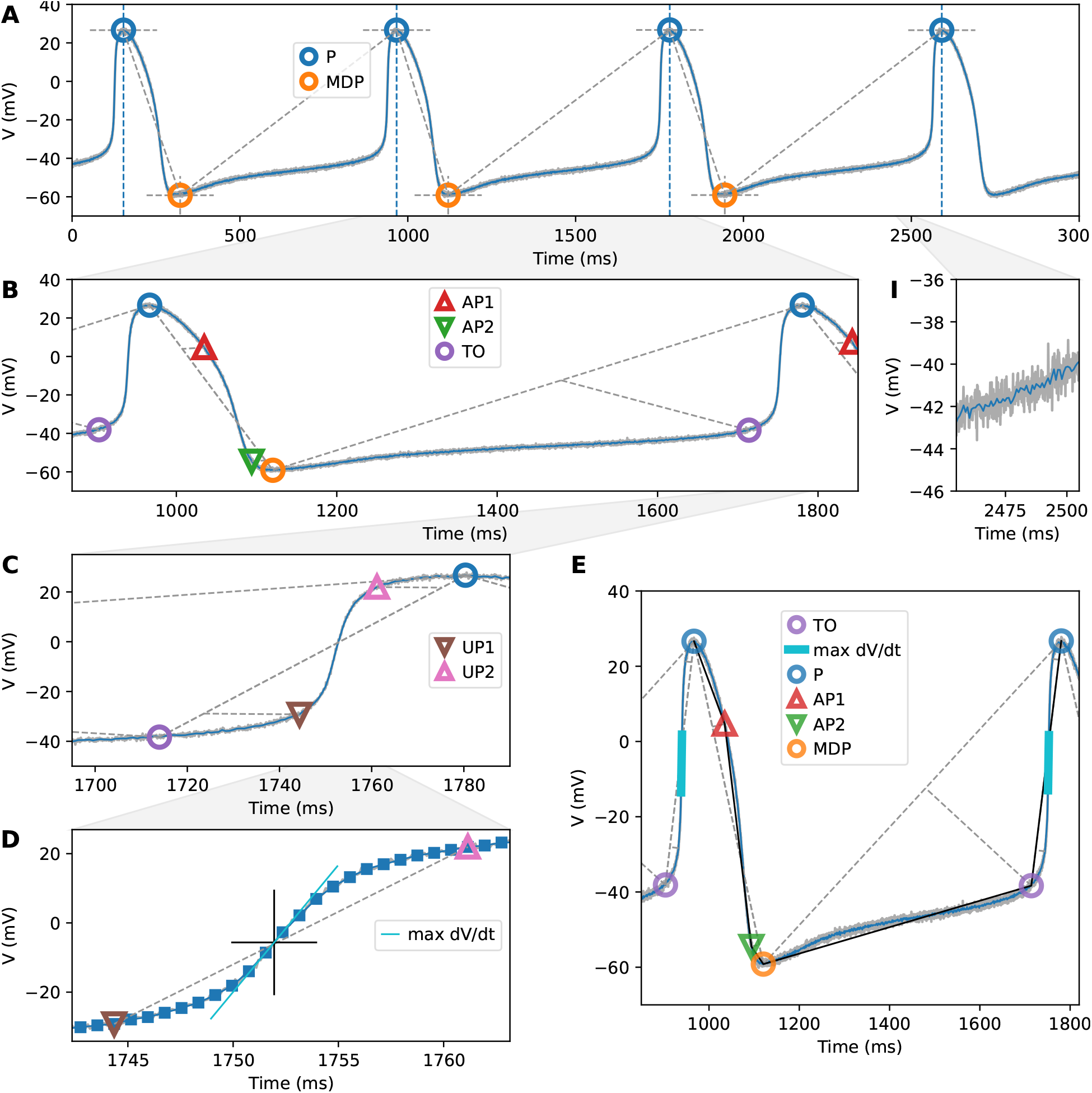
A linear approximation of a nodal AP. **A** In the first step, peaks (P) and MDPs are found by looking for regions where the voltage *V* is continuously above (resp. below) a threshold *v*_*t*_ for more than *n* samples, and then finding the maximum (resp. minimum) on those segments. The analysis is performed on a downsampled and smoothed trace (see panel **I**). **B** A line is drawn from peak to MDP, the signal is rotated so that this line is horizontal, and maximum/minimum along this line are denoted as AP1 and AP2. Similarly, the point furthest below the line from MDP to next peak is noted as TO (for “take-off”). Note how the dashed lines are perpendicular in the normalised space shown in panel E, where time and voltage are normalised by a characteristic scale), but do not look perpendicular in these panels. **C** In an attempt to characterise the upstroke speed, we zoom in on the section from TO_i_ to P_i+1_, and identify two new points, UP1 and UP2. However, these points still do not capture the steepest region of the upstroke. **D** Abandoning the geometrical methods, in this signal we find we can clearly identify the maximum upstroke velocity by inspecting the numerical first derivative of the downsampled signal. **E** The full linearisation. This panel is plotted with an aspect ratio equal to the ratio between our characteristic time and voltage scales, so that the perpendicularity of the dashed lines can be seen.

**Figure 2.**
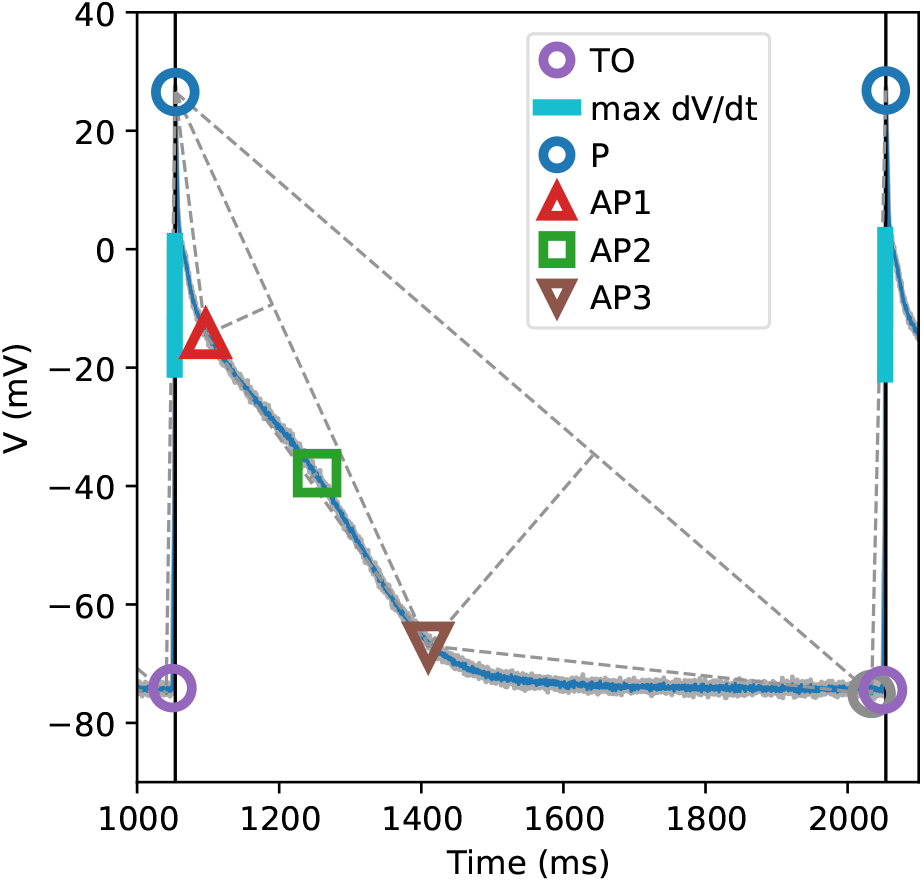
Points obtained on an atrial AP. The timing of the MDP point (indicated in gray) was determined by noise, so this point is left out of the final set.

**Figure 3.**
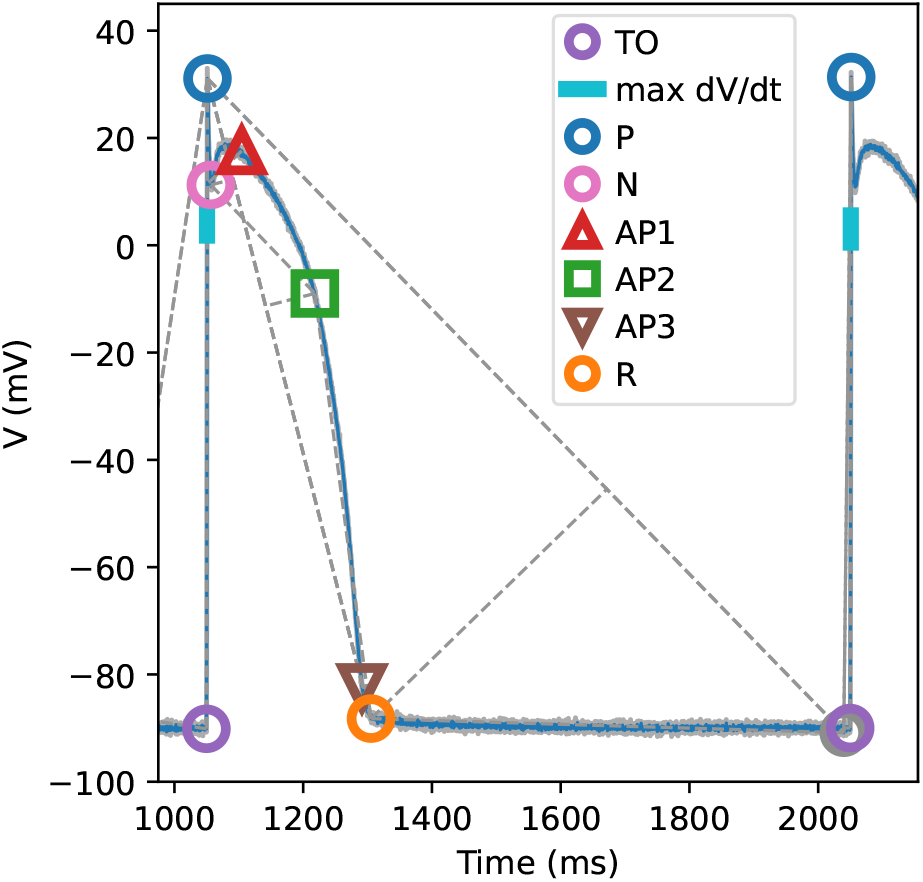
Points obtained on a ventricular AP. The MDP point is shown in gray, but mostly obscured by TO.

Using the same method more points could potentially be added. For example, a third point on the AP could be defined by finding the furthest point from the AP1-AP2 line, or the biphasic nature of the slow depolarisation leading up to the upstroke could be captured by adding a point from the line MDP-TO.

The area between TO_i_ and P_i+1_ proved problematic. Conventionally, this region is characterised by a physiologically important (but not always easy to measure) *maximum upstroke velocity*. As can be seen in Figure 1 *C*, the upstroke is not well characterised by the points TO and P, and adding further points (UP1 and UP2) did not improve the situation (Figure 1 *D*). Instead, a maximum upstroke velocity was found by non-geometrical means, by finding the largest sample-to-sample increase in V on the TO-P segment.

Next, we used the same approach to design biomarkers for an atrial prototype signal [5]. Here P and MDP were found as before, but the flatness of the diastolic region made the “MDP” location highly sensitive to noise. For this reason, we did not use the MDP in our final biomarkers, although we still use it in the construction of further points. Three points were defined to characterise the AP shape: AP3, perpendicular to the line P-MDP; AP1, perpendicular to P-AP3; and AP2, perpendicular to AP1-AP3. The point TO was defined as before. The upstroke was much faster than in the nodal AP, so that the maximum up-stroke velocity could not be obtained from the downsampled data — even after reducing the downsampling factor. Instead, the unfiltered data was used. However, this value might still be inaccurate if the 0.1ms sampling interval was insufficient.

We then applied our approach to a ventricular prototype signal [6, 7]. P and MDP were determined as before, and again MDP was sensitive to noise. As in the atrial case, we then added a point perpendicular to P-MDP, but in this case the rapid final repolarisation caused by strong I_K1_ meant that this point (R) indicated the end of the AP (rather than the APD_90_-analog found in the atrial case). Next, the points furthest from the line P-R were found and named N, for notch, and AP2. A point AP3 was found perpendicular to AP2-R, and AP1 was added perpendicular to N-AP1. This final point was intended to capture the plateau, but it may be better to simply take the maximum *V* on the segment N-AP2 instead. TO was obtained as before, and the upstroke was more rapid than even in the atrial case, making its velocity difficult to determine.

Finally, the table below shows how quantities similar to conventional biomarkers can be derived from the obtained points, using the atrial AP biomarkers from Saánchez et al.[1] as example. Similar quantities can be derived for the SAN and ventricular case.

| Atrial AP feature | Sánchez et al.[1] | Geometrical |
| --- | --- | --- |
| AP duration | $APD_{90}$ | $t_{AP3} - t_{TO}$ |
| Plateau duration | $APD_{50}$ | $t_{AP2} - t_{TO}$ |
| Spike duration? | $APD_{20}$ | $t_{AP1} - t_{TO}$ |
| AP amplitude | APA | $V_P - V_{TO}$ |
| Resting or minimum $V$ | RMP | $V_{TO}$ |
| Plateau potential | $V_{20}$ | $V_{AP1}$ |
| Upstroke velocity | $dV/dt_{max}$ | - |

## 4. Discussion

We showed how points characterising the shape of nodal, atrial, and ventricular APs can be obtained using a geometrical approach, and that these could be related to conventional biomarkers such as APD_90_. By “rotating” segments of the the signal and then finding maxima and minima, our method identifies points where the AP’s gradient changes, which may correspond to physiologically interesting points. This method could lead to more relevant points than ones using predetermined voltages (V_20_) or “percentages of repolarisation” (APD_20/50/90_).

Most atrial biomarkers defined in a previous study [1] could be redefined this way, with the exception of the maximum upstroke velocity, which had to be obtained in the conventional manner. However, it is interesting to note the sensitivity of this biomarker to data acquisition settings such as the sampling rate or any applied (hardware) filtering, so that although dV/dt_max_ is strongly correlated to important physiological quantities such as the magnitude of the sodium current and the conduction velocity, it may be worthwhile to explore alternative measures.

But are “geometrical” biomarkers more suitable for use in parameter estimation? From their derivation, we can expect that parameter changes causing small and smooth variations in the AP will lead to smooth changes in our biomarkers, resulting in a smooth error function. However, larger changes could cause points to “disappear” completely, leading to bifurcations in an error function that would be difficult to deal with. Future work is needed to test these ideas in practice, and evaluate the usefulness of geometrical versus conventional AP biomarkers.

